# Waning immunity drives respiratory virus evolution and reinfection

**DOI:** 10.1101/2024.07.23.604867

**Authors:** James J Bull, Katia Koelle, Rustom Antia

## Abstract

Reinfections with respiratory viruses such as influenza viruses and coronaviruses are thought to be driven by ongoing antigenic immune escape in the viral population. However, this does not explain why antigenic variation is frequently observed in these viruses relative to viruses such as measles that undergo systemic replication. Here, we suggest that the rapid rate of waning immunity in the respiratory tract is the key driver of antigenic evolution in respiratory viruses. Waning immunity results in hosts with immunity levels that protect against homologous reinfection but are insufficient to protect against infection with a heterologous, antigenically different strain. As such, when partially immune hosts are present at a high enough density, an immune escape variant can invade the viral population even though that variant cannot infect fully immune hosts. Invasion can occur even when the variant’s immune escape mutation incurs a fitness cost, and we expect the expanding mutant population will evolve compensatory mutations that mitigate this cost. Thus the mutant lineage should replace the wild-type, and as immunity to it builds, the process will repeat. Our model provides a new explanation for the pattern of successive emergence and replacement of antigenic variants that has been observed in many respiratory viruses. We discuss our model relative to others for understanding the drivers of antigenic evolution in these and other respiratory viruses.

## Introduction

Many respiratory viruses, such as influenza viruses, coronaviruses, and RSV, present challenges for control because individuals are repeatedly infected throughout life. Longitudinal cohort studies indicate that individuals can become reinfected with the same influenza virus subtype in months to years following a previous infection [35, 11]. Virological and serological studies on the four seasonal human coronaviruses also indicate that individuals are reinfected every few years [8, 12], and a recent systematic review indicates that similar patterns of reinfection are observed in SARS-CoV-2 [29]. RSV reinfection has also been shown to commonly occur [13]. Frequent reinfection is not characteristic of all human viruses. For example, we are very rarely re-infected with measles, yellow-fever, smallpox, mumps, rubella, or polio, all of which are viruses that undergo systemic replication.

The lack of reinfection observed for viruses with systemic replication is jointly due to the persistence of high levels of immunity and viral antigenic stability. Why, then, can respiratory viruses reinfect? It seems that the answer must lie either with waning host immunity or viral immune escape, or both (Fig 1). Antigenic changes in viral populations seem to be the obvious culprit underlying reinfection potential in respiratory viruses as these viruses commonly exhibit ongoing antigenic evolution [36, 31, 9, 20, 21, 39]. While antigenic changes are both associated with and contribute to reinfection, it is not clear why respiratory viruses typically exhibit antigenic evolution while viruses undergoing systemic replication do not.

**Figure 1:**
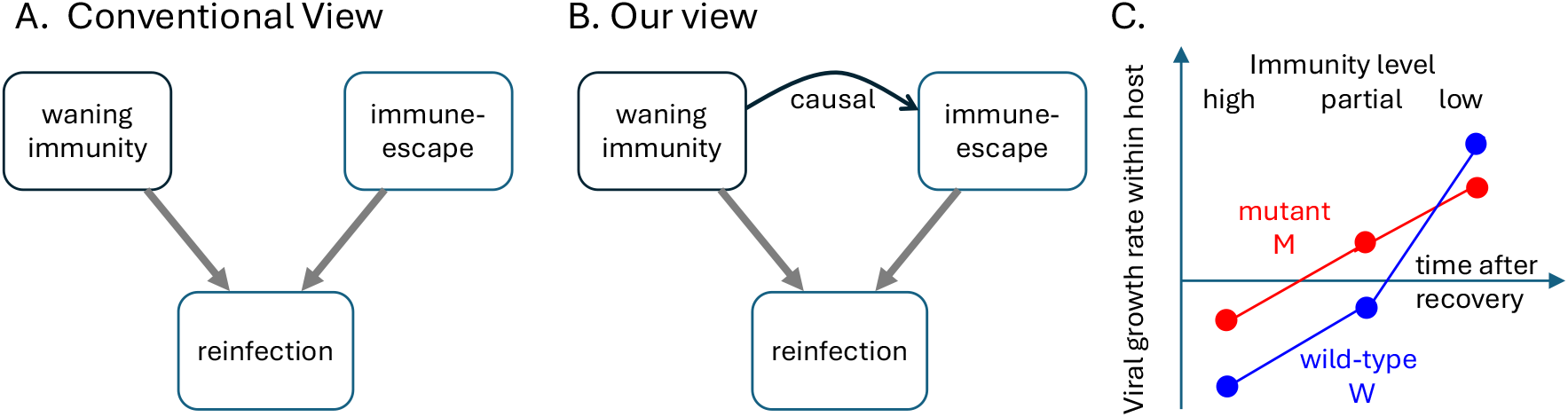
The relationship between waning immunity, immune escape, and reinfection in respiratory viruses. (A) A schematic depicting the conventional view of factors impacting host reinfection. Waning immunity refers to an individual’s decay in T-cell and antibody levels over time. Immune escape refers to genetic changes in the viral population that result in antigenic novelty. Here, an individual can become reinfected because their immunity has waned or because they are challenged with a strain that is antigenically different from their previous infection. (B) A schematic depicting our view of factors impacting host reinfection. Here, immune escape is not independent of waning immunity; rather waning immunity is a driver of immune escape in the viral population. (C) The impact of host immunity level on within-host viral growth rates that is assumed by our model. Following infection with a wild-type virus, individuals transition over time from having full to partial to low immunity levels. Neither virus can grow in (or transmit from) hosts with high levels of immunity to the wild-type virus. Both viruses can grow in (and therefore transmit from) hosts with low levels of immunity to the wild-type virus, but the mutant has a lower within-host growth rate due to the fitness cost of the antigenic mutation and thus has a transmission disadvantage in low-immunity populations. Only the mutant virus can grow in (and transmit from) hosts with partial levels of immunity to wild-type due to the antigenic difference between the mutant virus and the wild-type virus.

One possible explanation that has been invoked relies on variation in mutational tolerance across viruses: viruses that exhibit a greater degree of mutational tolerance may have more opportunity to evolve an immune escape variant that is still capable of efficient replication within a host. In support of this explanation, the hemagglutinin (HA) surface protein of influenza viruses shows greater mutational tolerance than the HA surface protein of the measles virus [19, 38]. A second explanation invokes variation in the breadth of the immune response across viruses: viruses that elicit a narrow response to an immunodominant epitope may more easily evolve immune escape than viruses that elicit a broad response against multiple epitopes [16]. This explanation is supported by neutralization studies in measles virus that indicate that antigenic sites are serologically co-dominant [28], while for influenza viruses, the immune response appears to be directed at only one to two immunodominant epitopes [43, 1]. While both of these explanations are plausible and each have some empirical support, it is not clear why factors such as mutational tolerance or the breadth of the immune response would differ systematically between respiratory viruses and viruses undergoing systemic replication.

Here, we propose an alternative explanation for why some viruses undergo antigenic evolution while others do not. Our explanation is rooted in known differences between the immune response to respiratory viruses versus to viruses undergoing systemic replication, and thus provides a more general explanation for observed differences between these two classes of viruses. In brief, immune responses to respiratory infections differ both qualitatively (antibody isotype and T cell phenotype) and quantitatively (in their rate of waning) from those of systemic infections. This is because the response to respiratory viruses must cross from the blood into the respiratory tract. Resident memory T cells, and antibodies of the IgA and IgG1 isotypes are actively transported from the blood to to the respiratory tract where they control viral replication [32, 6]. Subsequent to clearance of the infection, high titers of antibodies and large populations of B and T cells are maintained systemically for many decades following infection or vaccination [2, 3]. In contrast to systemic immunity which is long-lasting, immunity in the respiratory tract wanes rapidly with a half-life of weeks or months [24, 18, 23]. Consequently reinfections with respiratory viruses can be observed even when the virus does not change antigenically [11, 5, 14], whereas reinfection with viruses that replicate systemically is rare [41]. Our model proposes that this waning of immunity not only itself allows for reinfection, but that it is further a key driver of antigenic immune escape (Fig 1B), which in turn, exacerbates reinfection potential and is observed continually in respiratory virus populations at the level of the host population. Our model differs from an earlier model proposed by Grenfell et al [17] insofar as they suggested waning immunity facilitates the generation of escape variants at the within-host level while we focus on the spread of escape-variants in the population.

## Results

### Waning immunity drives viral evolution: intuition

At the host population level, waning immunity results in the generation of hosts with intermediate levels of immunity. The presence of these partially immune hosts can facilitate the emergence of viral immune escape in two distinct ways. The first is that immune escape variants may be produced at higher rates in partially immune individuals than in other infected individuals. This argument posits that the strength of the immune response in partially immune individuals is sufficiently low to allow for viral replication and thus the within-host generation of de novo immune escape variants, while it is also sufficiently high to provide these immune escape variants with a within-host selective advantage over the infecting virus [17, 25]. As such, the population-level rate at which immune escape variants are generated is effectively higher when the population comprises many partially immune individuals. This explanation therefore assumes that the emergence of antigenic escape variants is mutation-limited at the level of the host population.

A second, and yet unexplored possibility is that waning immunity creates the situation shown in Fig 1C, which depicts the within-host growth rate of two virus strains: the strain an individual has previously been infected with (hearafter, ‘wild-type’) and a newly circulating antigenic escape strain (hereafter, ‘mutant’). Within-host growth rates are shown for individuals under three different levels of immunity (high, partial, and low), which individuals pass through following infection. Individuals having been infected recently have high levels of immunity and within-host viral growth rates therefore fall below zero for both the wild-type and the mutant virus. As such, these individuals are not susceptible to infection with either the wild-type strain or the mutant strain. In contrast, individuals having been infected a long time ago have only low levels of immunity and within-host viral growth rates therefore fall above zero for both the wild-type and the mutant virus. These individuals are susceptible to infection with the wild-type strain and the mutant strain and would be able to transmit these infections to others. We assume that the mutant virus carries a replicative fitness cost and thus has a lower within-host viral growth rate than the wild-type virus in individuals with only low levels of immunity [37]. This leads to lower transmission potential of the mutant strain relative to the wild-type strain in populations of host with low levels of immunity. Individuals having been infected an intermediate length of time ago have partial levels of immunity, resulting in the within-host viral growth rate for the wild-type virus still falling below zero and that of the mutant virus exceeding one. These individuals are thus still not susceptible to infection with the wild-type virus but are susceptible to infection with the mutant virus. Waning immunity in this case has the potential to generate a class of partially immune individuals that could provide a selective advantage to the mutant virus at the level of the population, even if this virus harbors a within-host replicative fitness cost. This explanation therefore assumes that the emergence of antigenic escape variants is not mutation-limited at the level of the host population, but rather is selection-limited.

While the mutation-limited and the selection limited models are not mutually exclusive, they are quite distinct: the first argues that partially immune individuals increase the probability of generation of a mutant virus at the within-host level [17], while the second argues that partially immune individuals facilitate population-level transmission of the mutant virus. We argue here that this second possibility is the primary driver of antigenic evolution for respiratory viruses. As such, we propose that waning of immunity in respiratory viruses and immune escape are not two independent factors that enable host reinfection, but rather that waning immunity also drives immune escape (Figure 1B), placing waning immunity as the ultimately responsible culprit for respiratory virus reinfection. Our argument is a special case of the widely acknowledged principle for chemicals applied to suppress populations: intermediate levels of suppression foster the evolution of complete resistance in the long-run by enabling evolution of resistance in small steps. In the present context, an abundance of hosts with partial immunity provides a subpopulation that enables growth of mutant viruses that partially escape immunity. Those mutants can be thought of as ‘small-step’ mutants because they could not spread in a population of just susceptible and immune hosts. In this model, the introduction of small-step viral mutants is not limiting, but their ascent in the population requires a sufficient density of hosts with partial immunity.

Attaining a ‘sufficient’ abundance of partially immune hosts for a mutant to have a selective advantage rests on a combination of viral and host factors. Intuition suggests that, for the mutant to be able to invade, waning immunity rates should be neither too high nor too low. If immunity wanes too slowly, most recovered hosts will remain in the fully immune class, creating a shortage of partially immune hosts. If immunity wanes too quickly, recovered hosts will rapidly return to naïve status, where the wild-type has the transmission advantage. However, the basic reproduction number *R*_0_ of the virus (a measure of transmission potential) is also likely to be important, as a low *R*_0_ will lead to few infections and thus few recovered hosts – which are the precursors to partially immune hosts.

### Waning immunity drives viral evolution: a mathematical model

To better understand this intuition, we introduce a simple epidemiological model with waning immunity (Figure 2, Table 1). We first limit the model to a single (wild-type) virus and account for individuals who are susceptible to infection (*S*), individuals who are infected with the wild-type virus (*I*_*W*_), individuals who have recently recovered and are fully immune to the virus (*R*), and individuals who are partially immune (*P*). Following infection, we let immunity wane gradually, with individuals passing in to and out of the partially immune host compartment at rates *γ*_1_ and *γ*_2_, respectively. We assume that only individuals in the *S* class are susceptible to infection with the wild-type virus. These individuals may never have been previously infected or they may have had a previous infection a long time ago (such that they have transitioned from *R* through *P* and into *S* already. We assume individuals in the *P* class are not susceptible to infection with the wild-type virus. With variables *S, I*_*W*_, *R*, and *P* representing densities, the equations of the model are:

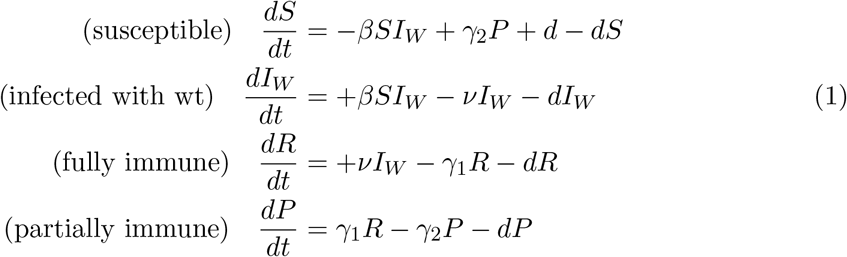

**Table 1:** Table of the variables and parameters. We use parameter values that we think are not biologically unreasonable.

| Symbol | Quantity | Value |
| --- | --- | --- |
| $S$ | Susceptible | |
| $I_W$ | Infected with wild-type virus | |
| $R$ | Fully immune | |
| $P$ | Partially immune | |
| $I_M$ | Infected with mutant virus | |
| $R_0$ | basic reproduction number of wt virus | varies |
| $\beta$ | transmissibility of wt virus | $R_0(d + \nu)$ |
| $1/\nu$ | duration of infection | 1 week |
| $1/\gamma_1$ | duration of immunity to mutant | varies |
| $(1/\gamma_1) + (1/\gamma_2)$ | duration of immunity to wild-type | varies |
| $1/d$ | lifespan | 50 yrs<sup>-1</sup> |
| $c$ | initial fitness cost to mutant | 0.2 |

**Figure 2:**
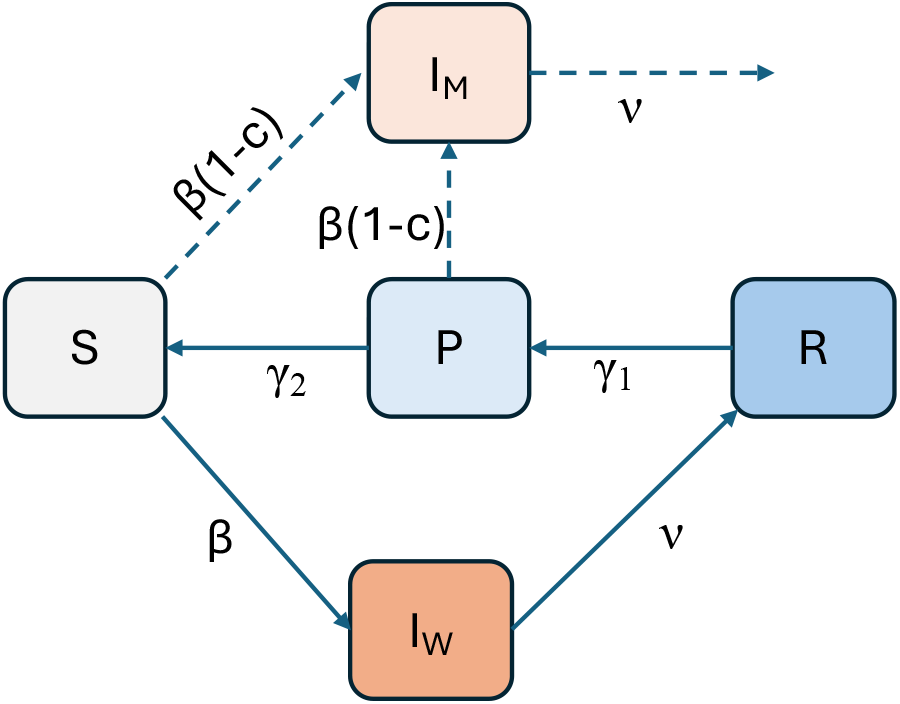
A schematic of the epidemiological model. Hosts are classified as susceptible to infection with the wild-type virus (*S*), currently infected with the wild-type virus (*I*_*W*_), fully immune (*R*), and partially immune (*P*). Individuals infected with the mutant virus are shown in compartment *I*_*M*_, although they are not explicitly modeled. The solid lines describe transitions involving infections with the wild-type virus. Immunity following infection wanes, with hosts first being fully immune (*R*) and not susceptible to infection with either the wild-type or the mutant virus. Hosts then transition to having partial immunity (*P*), which still protects them from wild-type infection but not from infection with the mutant strain. Dotted lines show transitions just after introduction of the mutant virus.

Individuals enter the susceptible class through birth (*d*) and through the loss of partial immunity (*γ*_2_*P*) and leave this class through infection (*βSI*_*W*_) and through background mortality (*dS*). Individuals enter the wild-type infected class through infection (*βSI*_*W*_) and leave this class through recovery (*νI*_*W*_) and background mortality (*dI*_*W*_). Individuals enter the fully immune class through recovery (*νI*_*W*_) and leave this class as their full immunity status wanes (*γ*_1_*R*) and background mortality (*dR*). Finally, Individuals enter the partially immune class through waning of full immunity (*γ*_1_*R*) and leave this class as their partial immunity status wanes (*γ*_2_*P*) and background mortality (*dP*). Given this model structure, the average duration of immunity to the wild-type strain is given by 1/*γ*_1_ + 1/*γ*_2_. The basic reproduction number (*R*_0_) for the wild-type virus in this model is given by *R*_0_ = *β*/(*d* + *ν*).

If *R*_0_ < 1, there is a trivial equilibrium with only susceptible individuals. If *R*_0_ > 1, there is an equilibrium with non-zero values for all classes 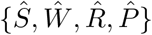 (see Appendix).

To consider the evolution of immune escape, we introduce infections with a mutant virus *I*_*M*_ into a population with the wild-type infection at equilibrium. The mutant virus differs from wild-type virus in that it has a reduced transmission rate, given by *β*(1 ™ *c*), where 0 *< c <* 1. Being antigenically different from the wild-type virus, this mutant can infect hosts with partial immunity. The dynamics of the mutant virus are thus initially given by:

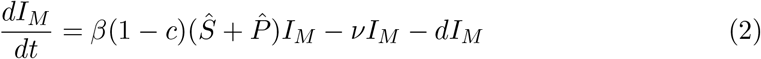

The mutant virus can invade at the level of the host population if its growth rate *r*_*M*_, given by 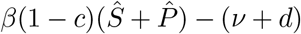, exceeds zero. Equivalently, the mutant virus an invade if its effective reproduction number, *R*_*M*_, exceeds one. *R*_*M*_ is in given by:

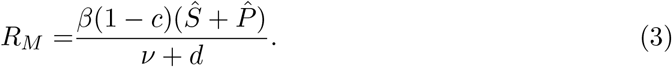

Because the equilibrium number of susceptible individuals in the population is given by *Ŝ* = 1*/R*_0_ (see Appendix), this expression can be simplified to:

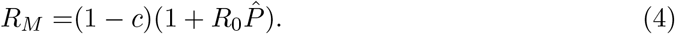

From this equation, we can see that the invasion potential of the mutant strain is entirely determined by its fitness cost *c*, the wild-type basic reproduction number *R*_0_, and the equilibrium density of partially immune hosts 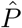. In Fig 3A, we first plot 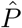 as a function of two parameters of the epidemiological model shown in Figure 2: the average duration of immunity to wild-type virus reinfection and the wild-type virus’s *R*_0_. The plot shows that the density of partially immune hosts is highest at high *R*_0_ and at an intermediate duration of immunity. As per equation (4), this should also be the region most favorable to invasion the mutant (that is, where *R*_*M*_ is highest). Indeed, when we plot *R*_*M*_ as a function of these same two parameters, under two different fitness costs (*c* = 0.2 in Fig 3B and *c* = 0.5 in Fig 3C), we see a broadly similar relationship between *R*_*M*_ and these underlying parameters. In both panels, we find that there is a zone defined by values of *R*_0_ and duration of immunity below which the mutant cannot invade (blue regions). Due to the direct effect of *R*_0_ on *R*_*M*_, the impact of *R*_0_ on *R*_*M*_ becomes more pronounced, with the invasion potential of the mutant (defined by *R*_*M*_) increasing with the *R*_0_ of the infection. This agrees with the intuition described in the previous section. As is evident from equation 4, a higher fitness cost *c* results in a direct reduction in *R*_*M*_, such that the region of parameter space where the mutant can invade is smaller at higher costs (Fig 3C versus Fig 3B).

**Figure 3:**
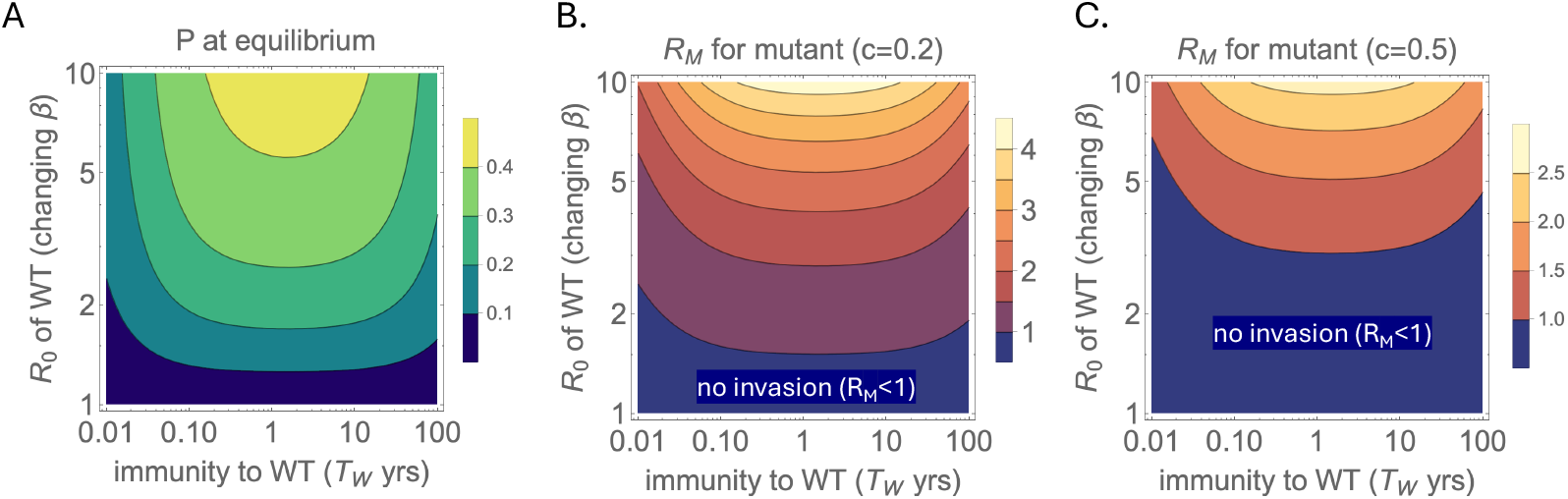
Invasion potential of the mutant virus that can infect partially immune hosts. (A) Equilibrium density of partially immune hosts 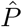 when just the wild-type virus is present. 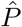 is shown as a function of two parameters: *R*_0_ and the average duration of immunity to reinfection with the wild-type virus. The latter is given by *T*_*W*_ = (1/*γ*_1_ + 1/*γ*_2_). Here, across all *T*_*W*_, we let the average duration of immunity to the mutant be half of that to the wild-type virus: 1/*γ*_1_ = (1*/*2)(1/*γ*_1_ + 1/*γ*_2_), which results in *γ*_1_ = *γ*_2_. (B,C) Calculated values of the mutant virus’s reproduction number *R*_*M*_ as a function of the wild-type virus’s *R*_0_ and the average duration of immunity to reinfection with the wild-type virus. In (B), *c* = 0.2. In (C), *c* = 0.5. Note that log scales are used on both axes in panels (A)-(C).

The invasion of the mutant depends on the relative time *T*_*M*_ = *γ*_1_ and and *T*_2_ = *γ*_2_ for immunity to wane from *R* → *P*, and *P* → *S* respectively. Fig 4 shows how invasion depends on the waning of immunity during the transitions from recovered hosts (*R*) versus partially immune (*P*) to susceptible (*S*). The advantage to the mutant is fast waning of ‘early’ immunity (small *T*_*M*_) and slow waning of ‘late’ immunity (*T*_2_). This asymmetry makes sense, because a small *T*_*M*_ quickly converts recovered hosts into hosts that can be infected only by the mutant, and a large *T*_2_ slows the rate at which those hosts again become susceptible to wild-type.

**Figure 4:**
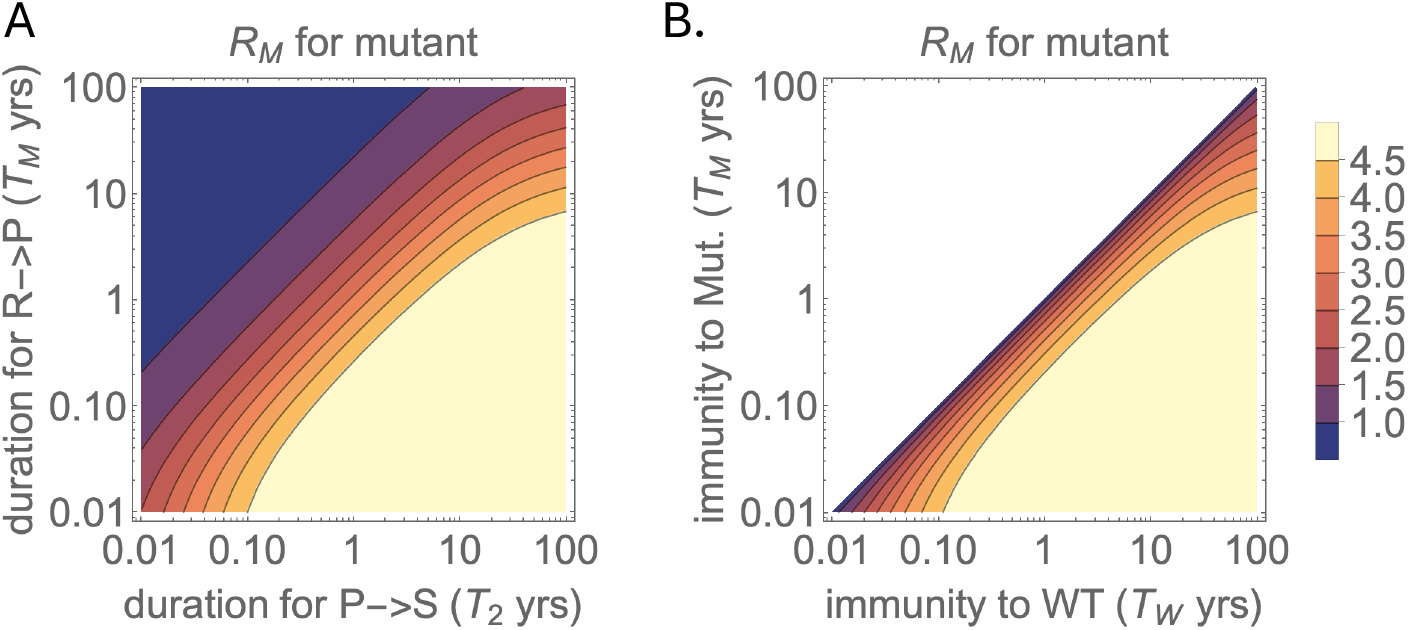
Mutant invasion benefits from different rates of waning to and from partially immune hosts. Parameter *T*_*M*_ = 1/*γ*_1_ is the rate for the *R* → *P* transition, *T*_2_ = 1/*γ*_2_ is for the *P* → *S* transition, and *T*_*W*_ = (1/*γ*_1_ + 1/*γ*_2_) is for the *R* → *S* transition. Both panels show in different ways that fast waning from *R* → *P* with slow waning of *P* → *S* favors the mutant. Panel (A) shows the effect of *T*_*M*_ and *T*_2_ separately. (B) shows the effect of *T*_*M*_ versus *T*_*W*_; the mutant invades more slowly as *T*_*W*_ approaches *T*_*M*_. Given that *T*_*W*_ is necessarily less than *T*_*W*_, the upper right is blank. Both panels assume *R*_0_ = 7 and cost, *c* = 0.2.

The model presented here merely considers the effect of waning immunity on the invasion of initial escape variants. We envisage that at the outset these escape mutations have only a small fitness advantage over the wild type virus, potentially because the antigenicescape is associated with a cost in terms of replicative fitness. However if *R*_*M*_ *>* 1, the mutant invades, and we expect that subsequent compensatory mutations will be rapidly generated and reduce this cost. The intrinsic growth rate of the mutant (in a population with no immunity) will then equal that of the wild-type virus [27, 40, 26, 15, 22]. Once the cost is mitigated, the mutant can replace the wild-type. We imagine this process will continue and can lead to successive replacement and increasing antigenic divergence from the original wild-type virus.

There are thus many elaborations to consider. We have also developed substantially more complicated models that incorporate measures of virulence and multiple stages of waned immunity; those models continue to support the role of waning immunity in favouring viral escape.

## Discussion

It is now well appreciated that immunity to many respiratory viruses wanes rapidly, and experimental studies suggest that individuals can be reinfected by the same strains more than once. We have suggested here that this waning immunity facilitates the evolution of viral escape from immunity in potentially small steps. The argument is merely that an abundance of hosts with partial immunity provides an environment that allows virus escape variants a transmission advantage over the wild-type when these escape variants could not spread in the absence of partial immunity. This argument will also hold for non-respiratory infections with waning immunity or when infection merely generates an intermediate level of immunity that does not wane over time.

The model presented here has been deliberately simplified to show the effect of waning immunity on mutant invasion. The primary reason for this simplicity is the lack of quantitative knowledge of the parameters governing both the waning of immunity, particularly in the respiratory tract, how it affects the dynamics of infection and transmission, as well as the costs and compensation associated with viral escape. Under these circumstances the simple model allows us to get an intuitive and robust understanding of the problem under investigation, namely the role of waning immunity for the emergence of antigenic variants. Furthermore the model identifies key features and parameters on how immunity wanes and on the costs and compensation associated with viral escape, features that warrant detailed investigation.

In the process depicted by our model, there is an interesting (albeit obvious) reciprocal interaction between waning immunity and evolution on viral infection. Waning immunity facilitates the evolution of antigenic escape which creates the appearance of faster waning of immunity (as depicted in Fig. 1B).

Our study focused on antigenic evolution that is observed over time at the population level for acute infections. Systemic virus such as HIV and HCV and protozoan infections such malaria and trypanosome infections can also exhibit antigenic changes. These changes frequently occur during the course of an infection and lead to longer lasting “persistent” infections. This process is different from what we have addressed.

We now turn to the relationship of our study to prior work. Grenfell et al [17] suggested that partly immune hosts are required for the within-host evolution of mutants that escape immunity. If these escape variants do not have a fitness cost, they will then have a selective advantage in the host population by being able to infect hosts with full immunity to the wild-type virus. A distinction between our model and theirs is that, in ours, the generation of mutants within a host is not limiting and the mutations initially are associated with a fitness cost. In our model escape mutations are assumed to be generated frequently (albeit with some initial fitness cost), and it is the widespread prevalence of partially immune hosts that allows – is required for – these escape-variants to invade. It is worth noting that within-host evolution of escape variants and the mechanism proposed here are not mutually exclusive: both can operate.

In the early days of covid vaccination, when vaccine supplies were limited, there were suggestions to halve the doses given to individuals so that more people could receive partial protection. This proposal was challenged on the grounds that weaker immunity (due to incomplete vaccination of individuals) would be more prone to select escape variants than would the immunity elicited by full vaccination [4, 33, 42]. The basis of this challenge was the same argument of [17] – within host selection of an escape variant. But that challenge was then met with a further counter argument, that weak immunity was not likely to select escape variants, instead that it was hosts with impaired immunity that were likely to be the source of escape variants [7].

Ferguson et al. [10] developed a model to explain the pattern of strain replacement that gives rise to the ladder-like phylogeny of viruses such as influenza A. They invoked a long-lived strain-specific immunity and a short-lived strain-transcending immunity. Our model suggests that this pattern of immunity arises naturally as a result of waning.

Alternative mechanisms for antigenic evolution – and for antigenic stability – have been proposed. We describe these by contrasting respiratory viruses such as influenza with the measles virus, which is so antigenically stable that the same vaccine has been effective for over 50 years. As mentioned by Yewdell [41], although measles is transmitted by the respiratory route, its replication occurs systemically, predominantly in B cells. Systemic immunity blocks systemic replication and prevents transmission even if pulmonary immunity has waned. From the perspective of our model, measles does not experience waning immunity, because once-infected hosts remain protected against transmission for life.

One alternative explanation for antigenic stability (not overlapping with ours) is merely that the proteins targeted by immunity are constrained from changing – the cost of immune escape is too high to evolve [38]. This hypothesis would seem to be refuted by the observed evolution of measles escape from monoclonal antibodies [34], but it remains possible that constraints are different *in vitro* than *in vivo*.

A more intriguing possibility has been suggested by [16]: measles virus fails to escape immunity because it elicits multiple codominant responses to several epitopes on the virus, while influenza has a single immunodominant epitope. We agree that multiple codominant responses (such as those to measles antigens) are required to prevent antigenic escape. However having multiple codominant responses is not sufficient if antibodies wane, as is likely to be the case for influenza. The waning of antibodies to all epitopes will eventually expose the virus to a selective regime where escape mutants to the most immunodominant response will be selected for escape.

Our models can be refined in a number of ways such as by considering the gradual waning of immunity (rather than a single class of partial immunity) and how this waning affects recall responses. Doing so will require a quantitative understanding of measures of immune efficacy that include susceptibility to infection as well as the level of transmission from infected individuals.

Our study has addressed a small problem in the grand picture of viral evolution and phylodynamics. Even if waning immunity proves to be a driver of viral antigenic evolution on a small, short term scale, there remain many puzzles in which this result should be integrated. Some of these have been alluded to in a preceding sections and include evolution beyond invasion and integrating the evolutionary and epidemiological scales to understand different patterns of antigenic evolution and phylodynamics of different respiratory infections. It will thus be important to understand whether our model is compatible with this broad scale pattern of evolution.

## Acknowledgments

We became aware that Bjarke Nielsen and Bryan Grenfell were developing an independent approach to a set of problems overlapping with ours. After consultation, we have opted to synchronize our submissions to bioRxiv [30]. We thank J. Yewdell for discussion.

## Appendix Steady States for text model 1

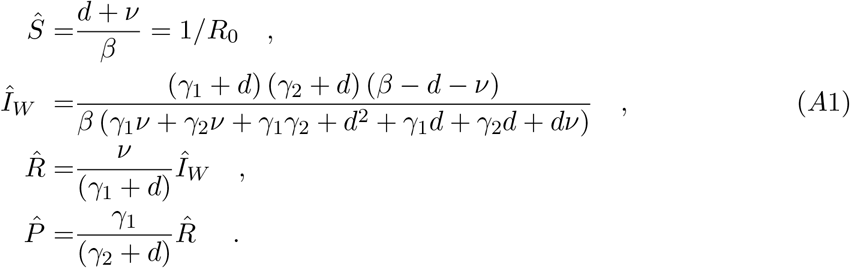

Notation is given in text Table 1.

